# Src-transformed cells hijack mitosis to extrude from the epithelium

**DOI:** 10.1101/246199

**Authors:** Katarzyna A. Anton, Mihoko Kajita, Rika Narumi, Yasuyuki Fujita, Masazumi Tada

## Abstract

At the initial stage of carcinogenesis single mutated cells appear within an epithelium. Mammalian *in vitro* experiments show that potentially cancerous cells undergo live apical extrusion from normal monolayers. However, the mechanism underlying this process *in vivo* remains poorly understood. Mosaic expression of the oncogene vSrc in a simple epithelium of the early zebrafish embryo results in apical extrusion of transformed cells. Here we find that during extrusion components of the cytokinetic ring are recruited to adherens junctions of transformed cells, stimulating formation of a misoriented pseudo cytokinetic ring. During extrusion, the ring constricts and separates the basal from the apical part of the cell releasing both from the epithelium. This process requires cell cycle progression and occurs immediately after vSrc-transformed cell enters mitosis. To achieve extrusion, vSrc coordinates cell cycle progression, junctional integrity, cell survival and apicobasal polarity. Without vSrc, modulating these cellular processes reconstitutes vSrc-like extrusion, confirming their sufficiency for this process.

---

At early stages of epithelial carcinogenesis, single mutations arise in single cells residing among normal epithelial neighbours within functioning organisms. In the past 10 years several laboratories around the world started uncovering a process called epithelial defence against cancer (EDAC)^1^. This is defined as a non-immunological primary defence mechanism whereby cells within an epithelial monolayer have the ability to sense a mutated neighbour and trigger pathways leading to its elimination. Although more recently the focus lied on the role non-transformed neighbours in EDAC^2–5^, there is evidence that transformed cells themselves have to undergo specific changes in the process of extrusion^6–10^. For example, in the case of vSrc-transformed cells (here referred to as vSrc cells), myosin activity regulated by myosin light chain kinase (MLCK) and Rho kinase (ROCK) as well as focal adhesion kinase (FAK) drive basal relocation of adherens junctions followed by apical extrusion^8^. Apart from mechanical shape adaptations, transformed cells residing among normal neighbours were also shown to undergo changes in basic cellular functions, leading to altered metabolism^9^ and endocytosis^10^. Until now, however, most studies of oncogenic cell extrusion have been performed using tissue culture models, cell lines and organoids, where cells are studied in environments different from the situation *in vivo*, such as matrix composed of just one protein, e.g. collagen I^7^, or glass^11^, a material of very high rigidity. These variable culturing conditions are known to affect cellular phenotype, behaviour and, especially, adhesion and cytoskeletal dynamics ^12,13^. Oncogenic cell extrusion requires a number of complex rearrangements within a fully differentiated epithelium, as well as determination of the direction of extrusion, as cells may either be extruded apically or basally^14^. Therefore, it is important to investigate this phenomenon within a living organism, where cells can extrude and delaminate freely.

Here, we performed a comprehensive mechanistic study of cell extrusion in living vertebrate embryos of zebrafish. Our model epithelium was the enveloping layer (EVL), the first polarised simple squamous epithelium that surrounds the yolk in the process of epiboly during gastrulation^15^. Unlike the *Drosophila* wing disc, the EVL is not pre-patterned in the anteroposterior and dorsoventral axes^16^, providing us with a homogenous cell population to study extrusion. Using the EVL-specific promoter Krt18, we established a system in which the tamoxifen-inducible transcriptional activator Gal4 (KalTA4-ERT2) was expressed exclusively within the EVL of the early embryo^3,10^ (Fig. 1A). In order to obtain mosaic inducible expression of a given oncogene, imitating early stages of carcinogenesis, we transiently injected constructs encoding oncogenes under the control of a UAS or double UAS (dUAS driving bi-directional expression) element at one cell stage. We have also created a double Krt18 promoter (dKrt18; Fig. S1A, B) resulting in constitutive expression of additional modulators of extrusion within the EVL. Thus, this *in vivo* system is reminiscent of *in vitro* models and the process of *in vivo* carcinogenesis, allowing us to generate two discrete cell populations: transformed and normal cells in a differentiated homogenous tissue.

Using this model, we document details of vSrc cell extrusion in the zebrafish EVL based on high-resolution live imaging. This approach uncovered a novel mode of extrusion in which vSrc holds the cell in the G2 phase of the cell cycle until a misoriented pseudo cytokinetic ring is formed and constricted in early mitosis, resulting in the cell leaving the epithelium.

## Results

### vSrc-transformed cells are apicobasally extruded from the zebrafish embryonic epithelium in a cell-death independent manner

We previously showed that when the oncogene vSrc was mosaically expressed in the EVL, transformed cells were apically extruded from the epithelium (outside of the embryo)^8^ (Fig. 1B, Movie 1). Further careful investigation of this process through live imaging revealed that transformed cells round up (become taller than their flat epithelial neighbours) and split into two fragments undergoing both apical and basal rather than exclusively apical extrusion (Fig. 1C, Movie 2). Apical parts were always larger than basal parts of extruding cells. The size of basal parts released towards deep cells of the embryo varied from at least a third of the total cell volume prior to extrusion (62% of vSrc-stimulated cell extrusion events) to smaller basal vesicles (38% extrusion events; data collected from 7 movies, 19 extrusions). Since apoptosis could result in cell fragmentation^17^, we investigated whether transformed cells died prior to becoming extruded. vSrc cells were found to be negative for cleaved-Caspase-3 prior and following extrusion (Fig. S2A). In contrast, EVL cells expressing death associated kinase 1 (DAPK1) died prior to becoming basally extruded (Fig. S2B). Moreover, inhibiting apoptosis by expressing the anti-apoptotic protein XIAP^18^ alongside vSrc, did not affect cell extrusion (Fig. S2C), although following extrusion, a larger apical fraction of vSrc-transformed cells eventually died, most likely via anoikis (Fig. 1C, Movie 2). Therefore, we concluded that vSrc-mediated extrusion was not due to cell death.

**Fig. 1.**
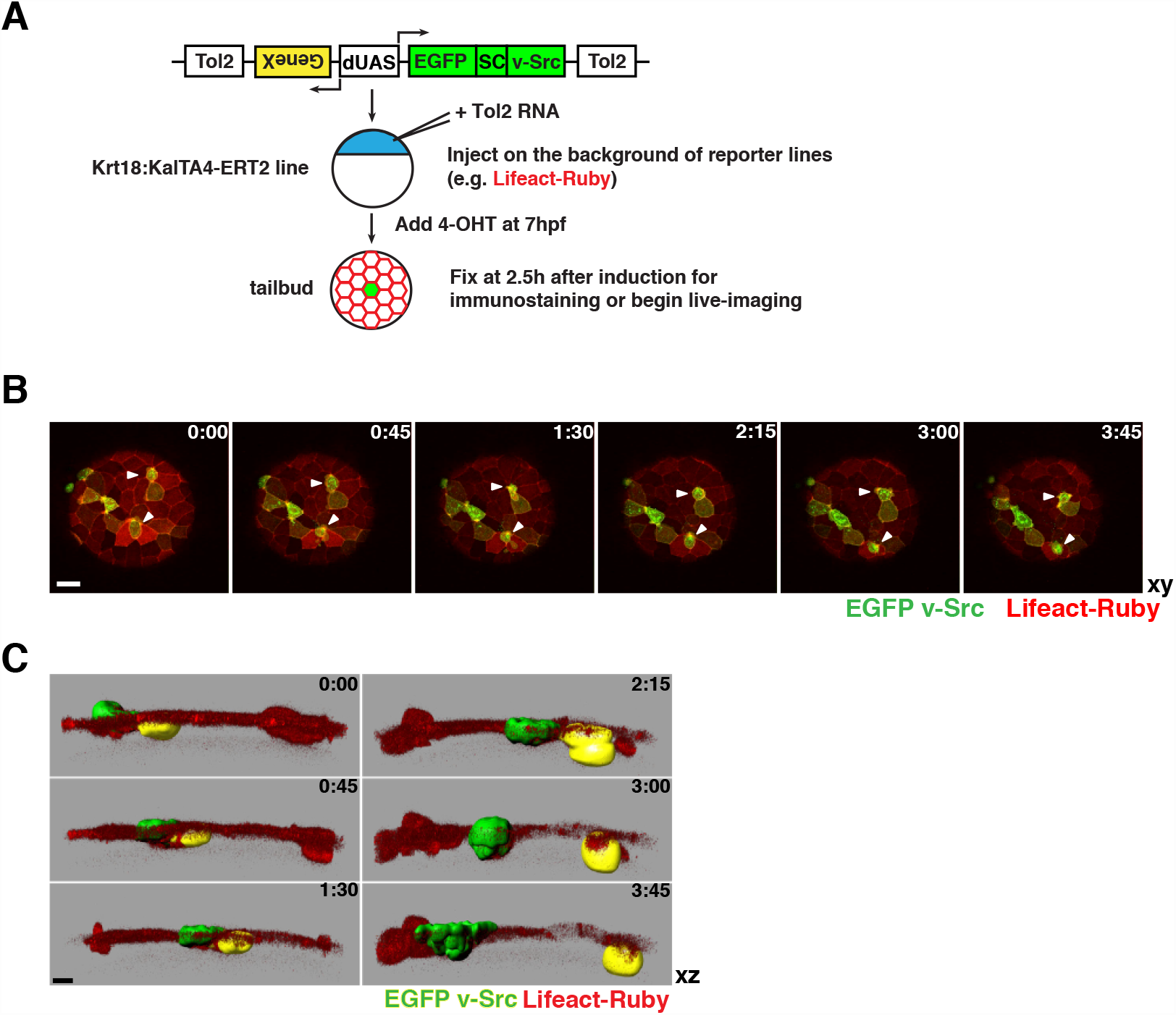
vSrc-transformed cells become apicobasally extruded from the EVL layer of the zebrafish embryo. **(A)** Experimental design. Fish embryos obtained from a transgenic line expressing tamoxifen-inducible Gal4 specifically in the EVL (Krt18:KalTA4-ERT2) are injected at one cell stage with constructs encoding oncogenes and effectors/markers under the control of the bi-directional UAS, dUAS^10^. At 50–70% epiboly, embryos are treated with tamoxifen to induce oncogene expression. At tailbud (2–3 hours from induction, 10 hours post fertilization), embryos are fixed for quantification or mounted in agarose for live-imaging. **(B)** Time-lapse imaging of vSrc cell extrusion from the EVL of the zebrafish embryo. Transgenic embryos obtained from a line expressing an RFP-actin marker (red) specifically in the EVL (Krt18:Lifeact-Ruby) line crossed with the Krt18:KalTA4-ERT2 line were injected with the UAS:EGFP-vSrc construct (green). Movies were taken over 4 hours. Frames were extracted from a representative movie at indicated times from the tailbud stage (t=0). White arrowheads indicate cells that become extruded. Scale bar, 50 *μ*m. **(C)** Segmented time-lapse images of vSrc cell extrusion. The surface function was used to segment GFP-positive vSrc cells over time using the Imaris software. In this cross section of the embryo (xz view), a cell is undergoing an apicobasal split (apical part extruding outside of the embryo is marked with green shading and the basal part extruding towards the deep cells with yellow shading). Scale bar, 10 *μ*m.

### vSrc aberrantly regulates cytokinetic machinery during extrusion

To investigate how vSrc cells produced apical and basal parts during extrusion, we further analysed their properties. Live imaging revealed that proliferation rates of vSrc cells within the EVL were significantly lower than control EVL cells expressing GFP only (Fig. 2A), likely because transformed cells were undergoing extrusion rather than mitosis (Fig. 2B). We then speculated that extruding vSrc cell used a contractile Actomyosin ring, such as the one assembled during cytokinesis, for this separation. Visualising Actin and Myosin in vSrc-mediated extrusion was inconclusive, as both proteins are constitutively present at the cell cortex of epithelial cells accumulating junctionally in ring-like structures coupled to adherens junctions (AJs)^19^ (see Life-actin-RFP in Fig. 1B). To visualise a contractile Actomyosin ring, we focussed on Anillin, a scaffolding protein required for the assembly of the cytokinetic ring^20^. During cytokinesis, Anillin is recruited to the mitotic plane through active RhoA, which is localised there by the negative and positive signals from the mitotic spindle^21^-24. Anillin in turn recruits Myosin and Actin, orchestrating the assembly of a contractile ring^20^. Following constriction, Anillin localises to the midbody, and to the nucleus in the interphase^25^. When imaged in normal EVL cells, Anillin-GFP behaved as described^20,25,26^ (Fig. S3A, Movie 3). However, in vSrc-expressing cells Anillin appeared to be recruited initially to junctional foci in the lateral region and eventually to a forming junctional ring, in addition to its nuclear localisation (Fig. 2C, Movie 4). This ring constricted during vSrc cell extrusion and in the moment of apicobasal separation resembled the midbody, which was apparently inherited by the apical part of the extruding cell (Fig. 2C). Importantly, the junctional localisation of Anillin has been reported in some tissue culture systems and in *Xenopus* embryos, where it is supposedly involved in junctional maintenance^27,28^. However, we did not observe junctional Anillin in normal EVL cells, with the exception of extrusion in vSrc cells and briefly in mitosis following nuclear envelope breakdown (NEB) prior to its recruitment to the mitotic plane.

**Fig. 2.**
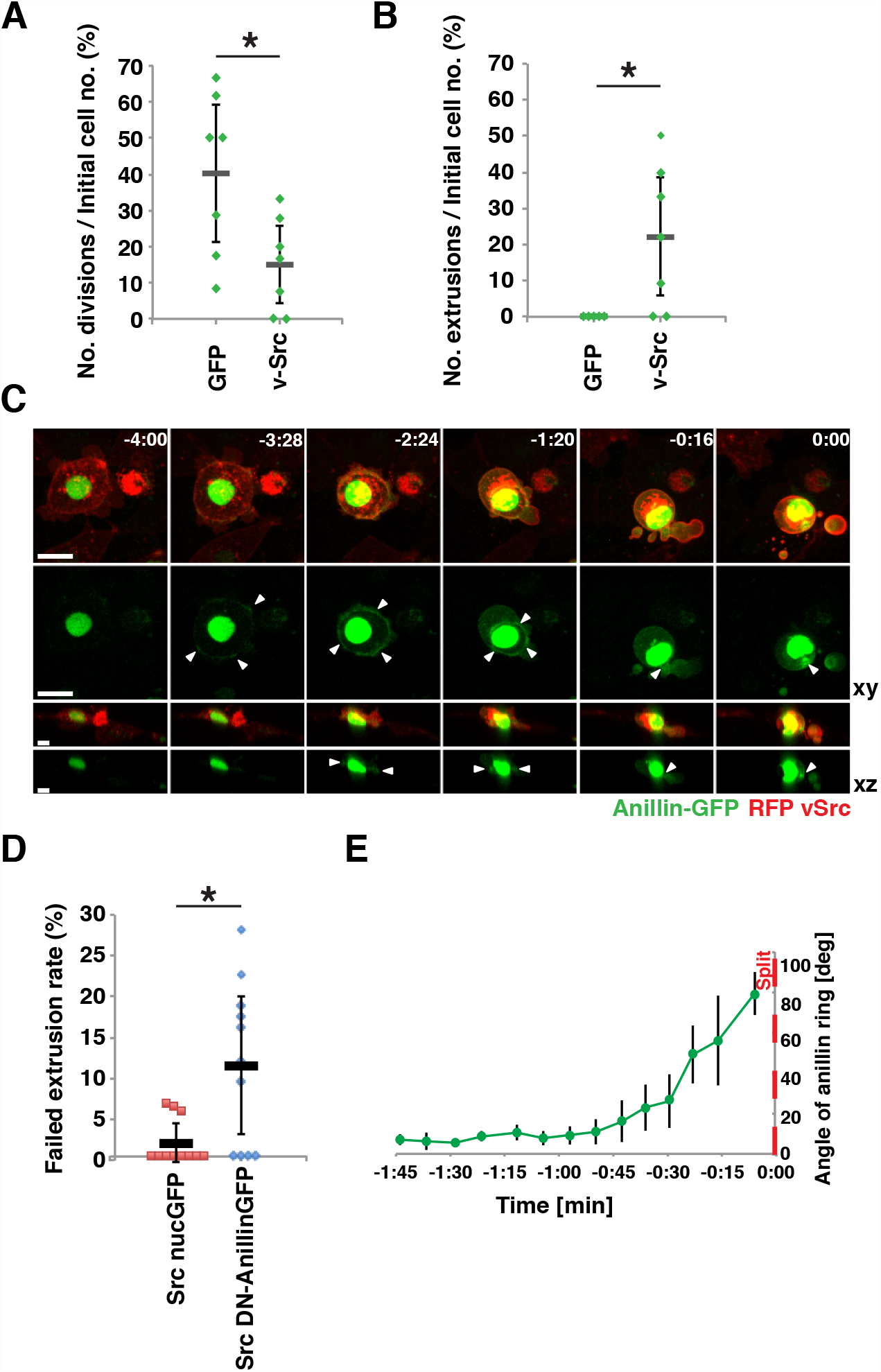
Contractile Anillin ring is recruited to the lateral cortex of vSrc cells prior to extrusion. **(A)** Quantification of EGFP and EGFP-vSrc cell division rates based on time-lapse imaging. Division rates were calculated as the number of divisions over 4 hours divided by the initial number of GFP-positive cells per embryo. Each mark represents a division rate in a single embryo. 7 embryos were imaged per condition in 14 independent experiments (total number of cells: n_GFP_ = 121, n_Src_ = 73). *P < 0.05. **(B)** Quantification of EGFP and EGFP-vSrc cell extrusion rates based on time-lapse imaging from (A). *P < 0.05. **(C)** Time-lapse imaging of Anillin-GFP during vSrc cell extrusion. Embryos were injected with the dUAS:myr-Cherry-vSrc;Anillin-GFP construct. Movies were taken over 4 hours. Frames were extracted from a representative movie at indicated times where t = 0 is the moment of extrusion. White arrowheads indicate the position of the Anillin ring. Scale bars, 25 *μ*m (xy) 10 *μ*m (xz). **(D)** Quantification of the failed extrusion rate based on time-lapse imaging. Embryos were injected with the following constructs: dUAS:EGFP-vSrc;nucGFP and dUAS:EGFP-vSrc;DN-Anillin. Failed extrusion rates were calculated as the number of cells that rounded up and then returned to the monolayer without division or extrusion over 4 hours by the initial number of GFP-positive cells per embryo. Each mark represents a division rate in a single embryo. Eleven embryos were imaged per condition in 3 independent experiments (total number of cells: n_Src_ = 168, nSrc,DNAnillin = 149). *P < 0.05. **(E)**Quantification of the angle between the Anillin ring and the surface of the embryo over time. Data from 4 cells acquired in 3 independent experiments were then aligned to the time of extrusion, t = 0, averaged ± s.d. and plotted.

The Anillin ring in vSrc cells appeared to be contractile during extrusion, as phosphomyosin light chain (pMLC) localised to the same plane (Fig. S3B). Additionally, we have created a dominant negative form of Anillin on the basis of a previously used mutated version in *C. elegans*^29^, which contains the ADH and PH domains (anillin and pleckstrin homology domains with the RhoA-binding motif) but lacks the MBD (myosin-binding) and ABD (actin-binding) domains. When expressed together with vSrc, this truncated form of Anillin significantly increased the rate of failed extrusions, during which cells become taller, round up and then are reintegrated into the monolayer, in comparison to vSrc alone (Fig. 2D). It is worth noting that, unlike most cultured cells^30^, EVL cells do not normally undergo mitotic rounding prior to mitosis (e.g. Fig S3A), therefore rounding-up indicates early stages of extrusion. Interestingly, the Anillin ring itself was initially parallel to the plane of the epithelium when a vSrc cell remained flat in the EVL. As the cell became taller and rounder during the extrusion process, the ring slowly shifted its orientation such that it was positioned more and more obliquely to the surface of the embryo towards the final moment of separation (Fig. 2E). Together, these data suggest that vSrc cells aberrantly utilise the cytokinetic machinery during extrusion.

### vSrc-driven extrusion is predominantly cell cycle-dependent and occurs at early mitosis

The ability of vSrc to modulate the cytokinetic machinery raises the question as to whether vSrc-driven extrusion requires mitotic entry. To investigate the role of cell cycle progression (Fig. 3A) in extrusion, we used fixed embryos, therefore all the following extrusion rates represent combined scores of “tall and apically extruded” cells as these remain attached to the surface of the embryo, while basally extruded parts move away and are impossible to account for. Expression of the G2/M transition inhibitors Wee1 (Fig. 3B) or constitutively active Pp2A (Fig. S4A) alongside vSrc led to strong inhibition of extrusion. Conversely, expression of a constitutively active form of the G2/M promoter Cdc25 (CA-Cdc25) resulted in an increase in extrusion (Fig. 3C). Wee1-mediated suppression of vSrc cell extrusion was rescued by constitutively active Cdk1 (Fig. 3D). Surprisingly, inhibiting G1/S transition did not have a significant effect on vSrc-driven extrusion (Fig. 3E). These results were subsequently confirmed when respective cell cycle arrests were achieved in the whole embryo with Emi1 abrogation^31^ (Fig. S4B, C; G2/M arrest) and a combination of the chemical inhibitors aphidicolin and hydroxyurea (Fig. S4D, E; G1/S arrest). These data suggest that there are two types of vSrc cell extrusion: a cell cycle-dependent mode for which a transformed cell has to enter mitosis and a cell cycle-independent extrusion that can occur in G1.

**Fig. 3.**
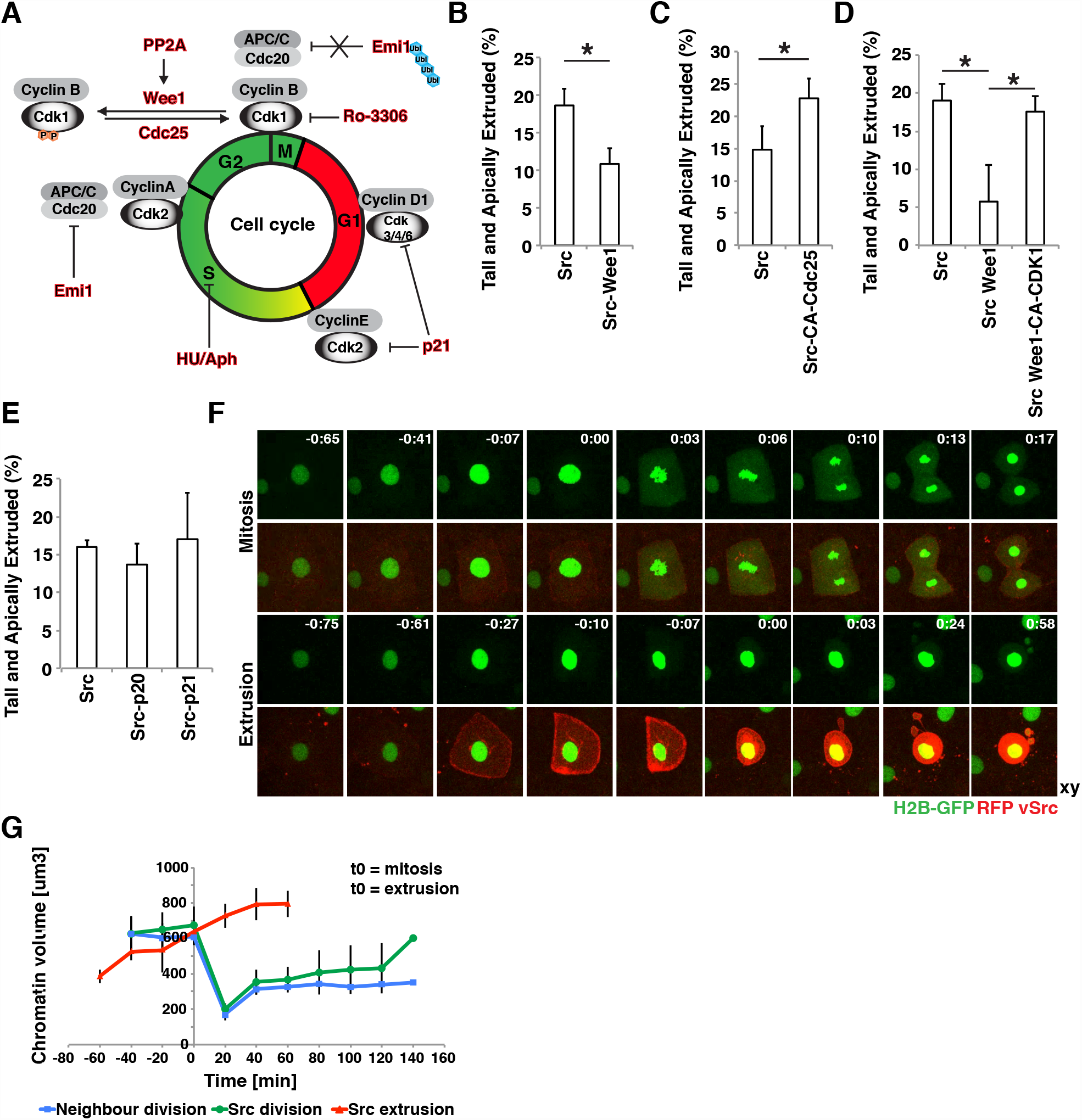
vSrc-transformed cells extrude in early mitosis, immediately after crossing the G2/M transition of the cell cycle. **(A)** A schematic model of cell cycle regulation. Highlighted in red are the molecules whose roles in cell extrusion were tested in this work. **(B)** The effect of Wee1 expression on vSrc-driven extrusion. Embryos were injected with the following constructs: dUAS:EGFP-vSrc and dUAS:EGFP-vSrc;Wee1. Data represent the number of tall and extruded cells divided by the total number of GFP-positive cells. Data are mean ± s.d. of 3 independent experiments (total number of embryos: n_Src_ = 21; nSrc,Wee1 = 27). *P < 0.05. **(C)**The effect of constitutively active Cdc25 on vSrc-driven extrusion. Embryos were injected with the following constructs: dUAS:EGFP-vSrc and dUAS:EGFP-vSrc;CA-Cdc25. Data are mean ± s.d. of 3 independent experiments (total number of embryos: n_Src_ = 18; n_Src,Cdc25_ = 21). *P < 0.05. **(D)** Constitutively active Cdk1 rescues Wee1 inhibition of vSrc-driven extrusion. Embryos were coinjected with the following constructs: dUAS:EGFP-vSrc alongside either dKrt18:myr-Cherry, dKrt18:Cherry-Wee1 or dKrt18:Cherry-Wee1;CA-Cdk1. Data are mean ± s.d. of 3 independent experiments (total number of embryos: n_Src_ = 30; nSrc,Wee1 = 29; nSrc,Wee1,Cdk1 = 33). *P < 0.05. **(E)** The effect of p20 and p21 expression on vSrc-driven extrusion. Embryos were injected with the following constructs: dUAS:EGFP-vSrc, dUAS:EGFP-vSrc;p20 and dUAS:EGFP-vSrc;p21. Data are mean ± s.d. 3 independent experiments (total number of embryos: n_Src_ = 21; nSrc,p20 = 19; nSrc,p21 = 11). **(F)** Time-lapse imaging of H2B-GFP in mitosis (top panel) and extrusion (bottom panel). Embryos were injected with the dUAS:myr-Cherry-vSrc;H2B-GFP construct. Movies were taken over 4 hours and frames were extracted from a representative movie. T0 indicates either the beginning of mitosis (chromatin condensation) or the moment of extrusion. Scale bars, 10 *μ*m. **(G)** Quantification of the chromatin volume (defined as the volume that H2B-GFP signal occupies in space) measured using the surface function of Imaris. The blue line follows chromatin volume change over time in a dividing EVL cell expressing myr-Cherry, the green line – in a dividing vSrc cell, the red line – in a extruding vSrc cell.

To establish whether extrusion is a form of misoriented mitosis, we imaged a number of mitotic markers in extruding vSrc cells. Visualisation of microtubules with Doublecortin (Dcx-GFP; Fig. S4F) and chromatin with Histone 2B (H2B-GFP; Fig. 3F) revealed that prior to extrusion the mitotic spindle is not assembled and full NEB does not occur. The nuclear volume is known to grow throughout the cell cycle and to reach its peak before mitosis^32^. Hence, we measured the chromatin volume in cells approaching mitosis or extrusion by determining the volume of the positive H2B-GFP signal (surface function of the Imaris software). We found that there was a set chromatin volume between 600–700 *μ*m^3^ at which EVL cells expressing myr-Cherry (membrane marker) or vSrc divide, and that the same volume was reached by vSrc cells immediately before extrusion (Fig. 3G). This observation indicates that vSrc cells reach the G2 phase and become extruded in early mitosis. Remarkably, the nucleus was always inherited by the larger apical part of the cell. We did not observe full NEB in extruding cells, although increased permeability of the nuclear envelope was detected; immediately before and after extrusion a portion of H2BGFP as well as nuclear GFP was present in the cytoplasm and inherited by the basal parts (Fig. 3F, Movies 5 and 6 and Fig. S4G). Overall, these observations indicate that most vSrc-transformed cells in the EVL become extruded in a cell cycle-dependent manner instead of completing mitosis.

### Src activation results in G2/M arrest that eventually leads to extrusion

To understand how exactly Src activation modulates cell cycle progression, we established a transgenic line expressing Cyclin B1-GFP (CcnB1-GFP) only in the EVL. Cyclin B1 forms a complex with CDK1 and allows entry into mitosis. Cyclin B1 is transcribed and stabilised in cells starting from the late S phase throughout the G2, and is rapidly degraded in mitosis^33,34^. Here we used it as a marker of the cell cycle phase in live imaging as none of the previously established FUCCI markers^35–38^ for imaging of the cell cycle in living embryos worked in our hands. In the CcnB1-GFP line, the GFP signal was present in the cytoplasm (from the late S phase) and gradually increased before mitosis. Following G2/M transition CcnB1-GFP localised first to the centrosomes, then to the nucleus^39,40^ and was finally degraded during division (Fig. 4A, Movie 7). To estimate the average length of different cell cycle phases in the EVL, we acquired movies of CcnB1-GFP transgenic embryos for the duration of 8 hours. Some of the cells that divided at the beginning of each movie divided again. For the quantification purpose, we split the cell cycle of EVL cells into 3 phases on the basis of changes in the intensity and localisation of CcnB1-GFP: (1) mitosis, defined as the time from the nuclear import of Cyclin B1 until completed cytokinesis, (2) G1/S phase, defined as the time from completed cytokinesis until the GFP signal reappeared in the cytoplasm, and (3) S/G2, defined as the time of the GFP signal present in the cytoplasm until its nuclear import. The average length of the cell cycle in the EVL was 8 h 25 min, with mitosis lasting 48 min and very variable G1/S and S/G2 lengths of on average 3 h 14 min and about 4 h 23 min, respectively (Fig. 4C).

**Fig. 4.**
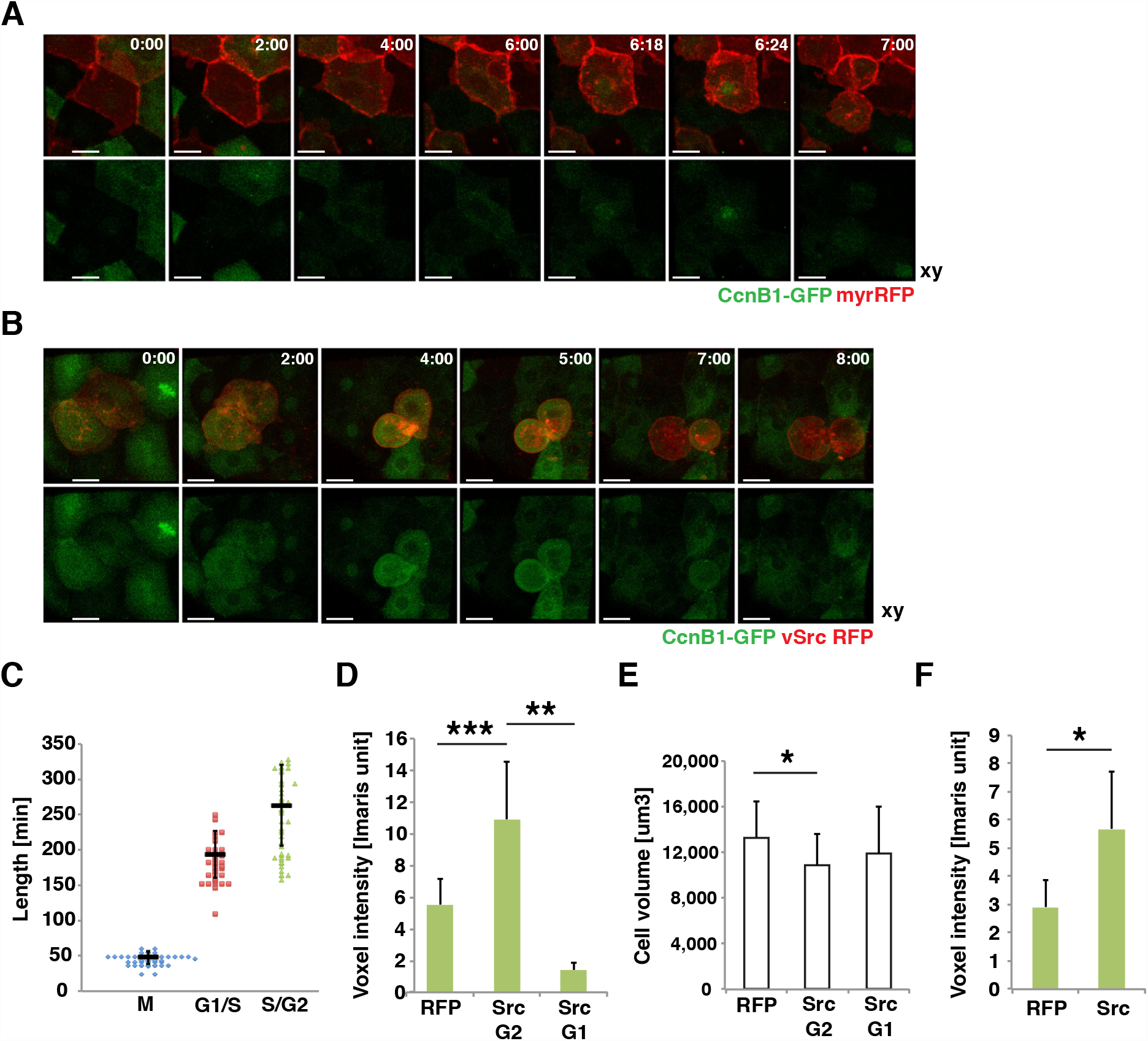
vSrc tyrosine kinase modulates the activities of major cell cycle players in the G2 phase prior to extrusion. **(A,B)**Time-lapse imaging of CyclinB1-GFP in cell extrusion. Transgenic embryos obtained from a line expressing a cell cycle progression marker specifically in the EVL (Krt18:CcnB1-GFP) crossed with the Krt18:KalTA4-ERT2 line were injected with the construct UAS:myr-Cherry (A) or UAS:myr-Cherry-vSrc (B). Movies were taken over 8 hours. Frames were extracted from a representative movie at indicated times from the tailbud stage. Scale bars, 10 *μ*m. **(C)** Length of different cell cycle phases of the EVL cells from the CcnB1-GFP line. The cell cycle phases are defined as follows; M is the time from the nuclear import of CcnB1-GFP until completed cytokinesis; G1/S is the time from completed cytokinesis until the GFP signal returns to the cytoplasm; S/G2/M is the time of the GFP signal present in the cytoplasm until mitosis. Each mark represents a single cell. Data collected from 5 movies in 3 independent experiments. **(D)** Mean voxel intensity ± s.d. of the CcnB1-GFP signal in single cells immediately before extrusion (myr-Cherry-vSrc) or division (myr-Cherry). Data collected from 31 extruded vSrc cells and 31 dividing myr-Cherry cells in 12 movies per condition from 5 independent experiments. ***P = 2.1×10^−7^, **P = 8.4×10^−4^. **(E)** Cell volume analysis before division and extrusion. Average volume ± s.d. of the cells used for CcnB1-GFP signal quantification immediately before extrusion or division (Fig. 4D). *P < 0.05. **(F)** Mean voxel intensity ± s.d. of the green channel (CcnB1-GFP) in myr-Cherry cells and myr-Cherry-vSrc cells at time 0 of a time-lapse from cells remaining in the epithelium. Data collected from 11 embryos per condition in 5 independent experiments. *P = 0.005.

To characterise how vSrc affects cell cycle parameters in the EVL, CcnB1-GFP transgenic embryos from the same batches were injected with constructs either carrying myr-Cherry-vSrc or myr-Cherry as a control. Each embryo pair (transformed and non-transformed) was then imaged and analysed side by side to avoid bias (representative pair: Fig. 4A, B, Movies 7 and 8). As a certain amount of the active Cyclin B1/CDK1 complex is required to trigger mitosis^34^, we measured the average intensity of the GFP signal in pairs of cells prior to division or extrusion. Firstly, 28 out of 31 extruded vSrc cells in CcnB1-GFP transgenic embryos had a high GFP signal, indicating that extrusion occurred in the later phases of the cell cycle (Fig. 4B). The average GFP intensity in vSrc cells about to be extruded was nearly doubled in comparison to equivalent control cells before division (Fig. 4D). Three out of 31 vSrc cells were extruded soon after dividing within the EVL in the G1 phase of the cell cycle, with very low average GFP intensity (Fig. 4D). The average volume of vSrc cells immediately before extrusion was significantly smaller than that of an average EVL cell before division, but this relatively small decrease was not sufficient to explain the average GFP intensity increase observed in vSrc cells (Fig. 4E). Moreover, the increased GFP signal could already be observed in vSrc cells remaining in the epithelium at the start of imaging (t = 0) as compared to control cells (Fig. 4F). Finally, when assessing how long vSrc cells remain in the S/G2 phase, we realised that cells, which eventually became extruded, were very rarely negative for the cytoplasmic GFP signal (only in 2 out of 28 cases), making it impossible for us to measure the length of the S/G2 phase prior to extrusion. Together, these data indicate that vSrc cells before extrusion remain longer in the S/G2 phase and accumulate more Cyclin B1 than normal EVL cells prior to division. This suggests that G2/M arrest presumably occurs prior to vSrc cell extrusion.

### Src-transformed mammalian cultured cells become extruded in cell cycle-dependent and cell cycle-independent fashions

To elucidate whether coordination of oncogenic extrusion and the cell cycle was a general phenomenon present also in mammals, we took advantage of the previously established tissue culture system utilising Madin-Darby Canine kidney (MDCK) mammalian cells^8^. Firstly, we treated mixed cultures of Src-expressing and wild type MDCK cells (mixing ratio 1:50) with cell cycle inhibitors: a CDK1 inhibitor Ro-3306 (G2/M arrest; Fig. S5A, B) and hydroxyurea (early S phase arrest; Fig. S5C). Consistent with the observation in the zebrafish EVL, we found that Src cell extrusion was significantly inhibited only by Ro-3306 (Fig. 5A). Further, we exploited the reliability of cell cycle markers for live imaging in mammalian cells and established a new line harbouring both active Src and the cell cycle monitor marker FUCCI^37^, which displays nuclear staining depending on the phase of the cell cycle: red in G1 and green in S/G2. We used this line for live imaging of Src cell extrusion (Fig. 5B) and found that 50% of Src-transformed cells in the mammalian system became extruded and subsequently their nucleus turned red (transition to G1 after extrusion; Fig. 5C). In some cases extrusion and division happened simultaneously, in others extrusion took place instead of mitosis (Fig. 5B). However, 30% of Src-transformed MDCK cells remained red throughout extrusion (stayed in G1 before and after extrusion, Fig. 5C), implying a cell cycle-independent mode of extrusion, similar to the one hypothesised in the embryo arrested in the G1 phase of the cell cycle. These data indicate that both the zebrafish embryo and mammalian cells utilise the two modes of oncogenic Src-driven extrusion, of which the more frequent one requires coordination with the cell cycle.

**Fig. 5.**
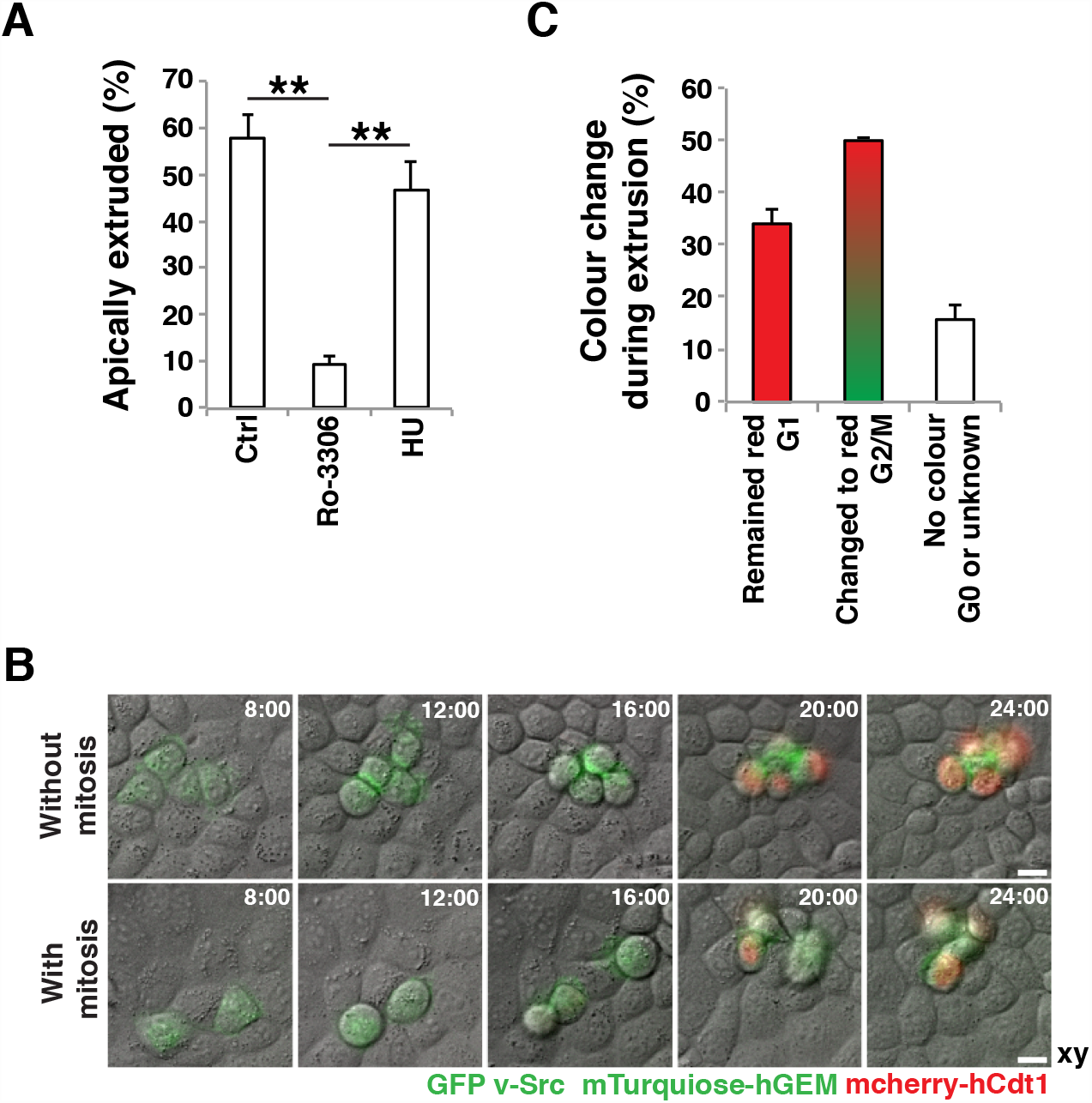
Src-transformed MDCK cells become extruded predominantly either during or instead of mitosis. **(A)**The effect of inhibition of the G2/M transition or G1/S transition on extrusion of MDCK pTR-cSrc-Y527F from a monolayer of normal MDCK cells. Inhibitors 2 mM hydroxyurea or 10 *μ*M Ro-3306 were added with a 9-hour delay after the induction of Src expression. At 24 hours from Src induction cells were fixed, stained with phalloidin and imaged. Data are mean ± s.d. 3 independent experiments (total number of cells: n_Ctrl_ = 155; n_HU_ = 153; n_Ro3306_ = 152). **P<0.005. **(B)** Time-lapse imaging over 24 hours of extrusion of MDCK stably expressing both pTR-cSrc-Y527F-GFP and FUCCI. GFP-CAAX labels Src cells (green membrane) that express Cherry-hCdt1 in G1 phase (red nuclei) or mTurquoise-hGEM in S/G2/M phases (green nuclei). **(C)** Quantification of the nuclear colour change in MDCK pTR-cSrc-Y527F-GFP/FUCCI cells during extrusion based on live imaging. Data are mean ± s.d. of 2 independent experiments (total number of extruded cells: n = 63).

### vSrc modulates cell cycle regulators Cdk1 and Pp2a

To investigate how vSrc modulates the G2/M transition, we searched for Src-phosphorylated proteins in the database (www.phophositeplus.org), and speculated that two good candidates for mediating this process are CDK1 and PP2A. PP2A is a phosphatase that antagonises the CDK1-CyclinB1 complex throughout the cell cycle, but has to be inactivated in order for a cell to enter mitosis^41^. Src phosphorylates PP2A on an inhibitory site tyrosine 307^42,43^, thereby promoting mitotic entry. As shown earlier, a mutant form of Pp2A, which lacks this phosphorylation, leads to an inhibition of vSrc-induced extrusion (Fig. S4A). Much less well documented is the action of the Src kinase on CDK1. The most probable Src-phosphorylation site within CDK1 is Tyrosine 15 (Y15), one of the two inhibitory sites that need to be dephosphorylated for activation of the mitotic complex CDK1-CyclinB1^44^ (Fig. 6A). It has been demonstrated that Src phosphorylates a CDK1 peptide surrounding Y15 *in vitro*, since this peptide has been used as a positive control for Src phosphorylation^43^. Therefore, we hypothesised that simultaneous phosphorylation of Pp2A and Cdk1 by vSrc, the former promoting, the later inhibiting G2/M transition, could be responsible for the prolonged G2 phase of vSrc cells. Hence, we sought to test that Cdk1 is a target of vSrc in our system.

**Fig. 6.**
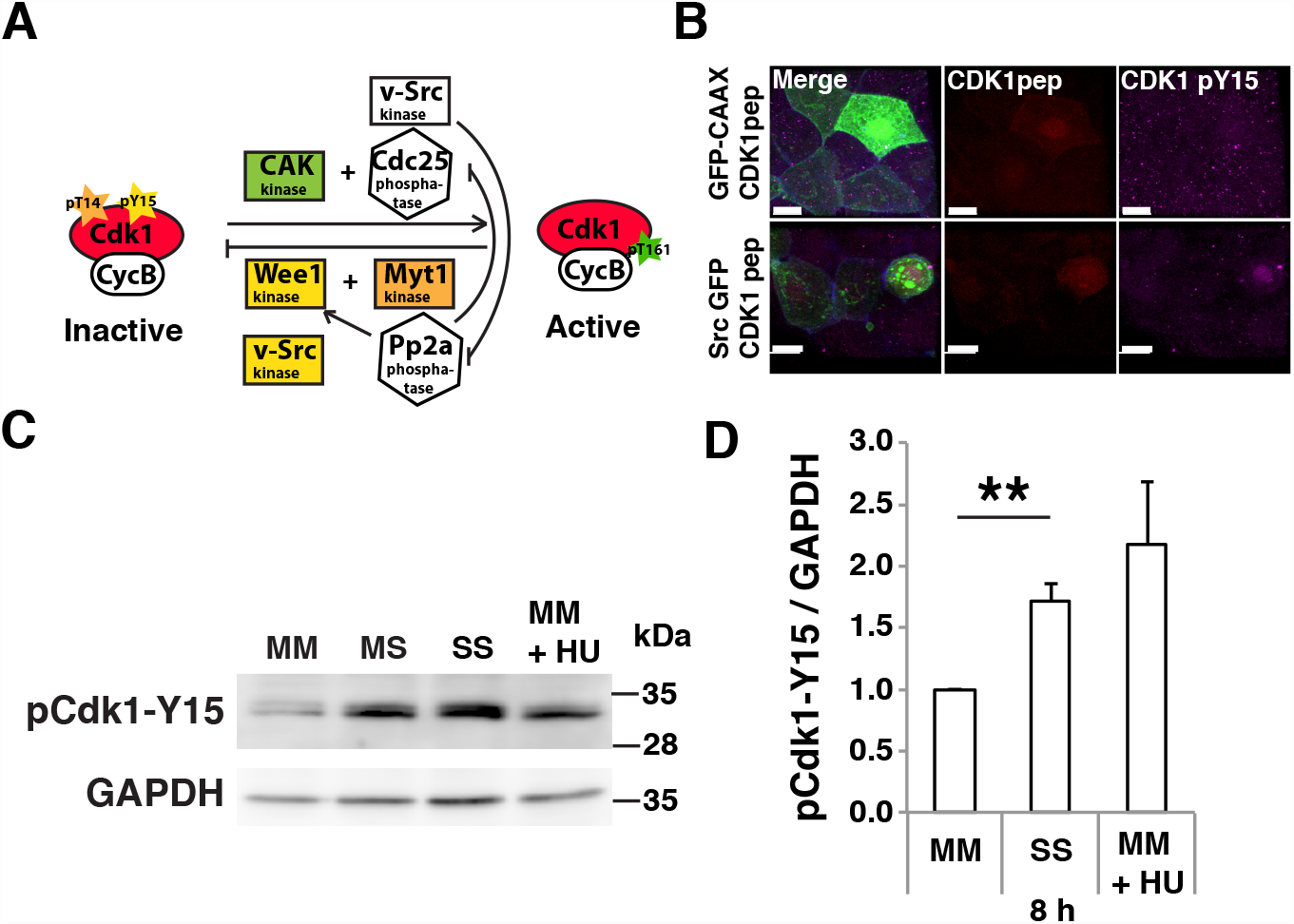
vSrc modulates cell cycle regulators CDK1 and Pp2a. **(A)** A schematic model of Src interference with cell cycle regulation. **(B)** Immunofluorescence images of CDK1 pY15 (purple) in the EVL cells expressing the mKO2-CDK1-pep (red) alongside EGFP or EGFP-vSrc (green). Scale bar, 10 *μ*m. **(C)** The effect on phosphorylation of CDK1 after 8 hours from Src activation in MDCK cells. MM – MDCK cells alone, SS – Src cells alone, MS – cultures mixed 1:1, MM+HU – MDCK cells treated with 2mM hydroxyurea (HU). **(D)** Quantification of the mean normalised signal ± s.d. in western blotting with the anti-CDK1-pY15 antibody after 8 hours from Src activation in MDCK cells from 3 independent experiments.

Since an anti-CDK1-Y15 antibody did not recognise zebrafish Cdk1-Y15 at endogenous levels, we created the authentic CDK1 peptide^43^ tagged with both RFP and an HA epitope. This peptide was phosphorylated prior to extrusion when expressed together with GFP-vSrc, but not with GFP-CAAX (Fig. 6B). To consolidate this observation, we turned to the MDCK tissue culture system. Normal MDCK cells alone, Src-expressing MDCK cells alone and mixed cultures were analysed by western blotting with the anti-CDK1-Y15 antibody. A low level of phospho-CDK1 was observed in normal MDCK cells, and the level was increased after incubation with hydroxyurea (G1/S arrest; Fig. 6C, D). The phospho-CDK1 level was much higher in Src-expressing cells and was moderate in the mixed cultures, suggesting that Src has the ability to directly or indirectly promote CDK1 phosphorylation, thereby inhibiting its activity (Fig. 6C, D).

A state in which cell cycle progression is both promoted and inhibited has been studied before and may result in “mitotic collapse”^45^. “Mitotic collapse” occurs after entry to mitosis, when CDK1 activation is not sustained at a high enough level for mitosis to proceed. This leads to dephosphorylation of mitotic substrates without degradation of Cyclin B1 and eventually results in cell death. Since Src activation both promotes and inhibits cell cycle progression, we wondered if cell extrusion could be a result of “mitotic collapse”. To recapitulate this state in the EVL without vSrc, we simultaneously expressed two G2/M modulators: the inhibitor kinase Wee1 and the active phosphatase CA-Cdc25 both directly regulating CDK1. Although we managed to block EVL cells at the G2/M transition (as confirmed by nuclear localisation of Cyclin B1 in Fig. S6A), no extrusion was observed at least over 4 hours. Inflicting other mitotic defects in the EVL such as triggering monopolar spindles, which can be induced by blocking Kif11 (Eg5) with an STLC inhibitor treatment^46,47^, also failed to cause extrusion (Fig. S6B, C). We concluded that EVL cells have the ability to cope with mitotic defects and delays without initiating death or extrusion of the “faulty cells” at least within the duration of our experimentations.

Overall, Src activation leads to a prolonged arrest in the G2 phase of the cell cycle due to modulation of cell cycle regulators Cdk1 and Pp2a. However, this modulation when mimicked in the EVL without vSrc, is insufficient to result in extrusion, suggesting that other vSrc effectors must be involved.

### vSrc modifies adherens junctions to recruit Anillin

At this point, the means by which vSrc hijacked the cytokinetic machinery were still unclear. In a dividing cell positioning of the mitotic plane is determined by the mitotic spindle^21,23^; however, the spindle and its cues for cytokinetic ring assembly were absent in a vSrc cell undergoing extrusion (Fig. S4F). This raised the possibility that the Anillin ring may be involved in extrusion via junctional constriction. Since RhoA activation promotes Anillin recruitment to the mitotic plane^20,22,24^ and modulates junctional integrity^48^, we investigated the effects of constitutively active and dominant negative RhoA on extrusion. Surprisingly, expression of either of these forms supressed vSrc-mediated increase in height (rounding-up), but not extrusion itself (Fig. 7A). Moreover, RhoA activation without vSrc did not trigger extrusion (Fig. S7A). These results imply that focal RhoA activation at the junctions is necessary for the assembly of the contractile Anillin ring, but that widespread activation or inactivation of RhoA presumably inhibits this process.

**Fig. 7.**
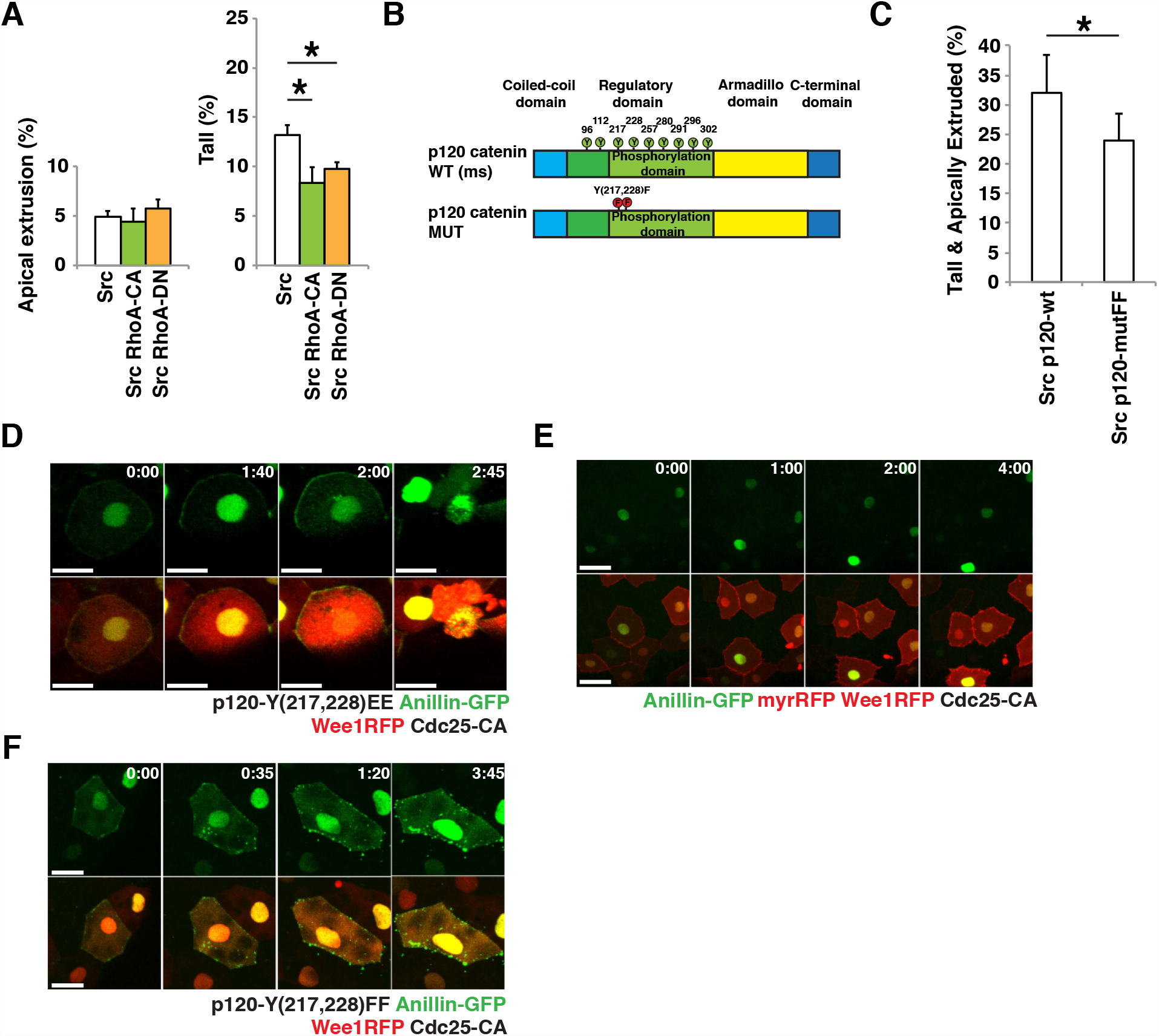
Src-phosphorylated p120-catenin recruits Anillin to the junctions allowing formation of a contractile ring. **(A)** The effect of constitutively active RhoA and dominant negative RhoA expression on vSrc-driven extrusion. Embryos were injected with the following constructs: dUAS:EGFP-vSrc, dUAS:EGFP-vSrc;CA-RhoA or dUAS:EGFP-vSrc;DN-RhoA. Data are presented in two graphs displaying cells “Apically extruded” (outside of the embryo) and “Tall” (remaining in the monolayer but taller and rounder than neighbours). Data are mean ± s.d. 3 independent experiments (total number of embryos: n_Src_ = 35; n_Src,CA-RhoA_ = 38; n_Src,DN-RhoA_ = 36). *P < 0.05. **(B)** A schematic model of the domain composition of p120-catenin. Highlighted in green are phosphorylation sites regulated by the Src kinase (from: PhosphoSitePlus database). In the bottom panel, a design of p120-catenin mutant with two sites, which regulate the interaction between p120-catenin and RhoA^51^. These Src-dependent phosphorylation sites are mutated from Y to F (p120-mutFF). **(C)** The effect of phospho-mimetic p120-mutFF on vSrc-driven extrusion. Embryos were injected with the following constructs: dUAS:EGFP-vSrc;p120-wt or dUAS:EGFP-vSrc;p120-mutFF. Data are mean ± s.d. 3 independent experiments (total number of embryos: n_Src_ = 34; n_Src,CA-RhoA_ = 42). *P < 0.05. **(D)** Time-lapse imaging of the effect of phospho-mimetic p120-mutEE on the localisation of Anillin-GFP in cells arrested at the G2/M transition. Embryos were injected with a combination of the following constructs: dUAS:Cherry-Wee1;CA-Cdc25 and dUAS:p120-mutEE;AnillinGFP. Movies were taken over 4 hours. Frames were extracted from a representative movie at indicated times from the tailbud stage. Scale bars, 25 *μ*m. **(E)** Time-lapse imaging of Anillin-GFP localisation in cells arrested at the G2/M transition. Embryos were injected with a combination of the following constructs: dUAS:Cherry-Wee1;CA-Cdc25 and dUAS:myr-Cherry;AnillinGFP. Movies were taken over 4 hours. Frames were extracted from a representative movie at indicated times from the tailbud stage. Scale bars, 50 *μ*m. **(F)** Time-lapse imaging of the effect of p120-mutFF on the localisation of Anillin-GFP in cells arrested at the G2/M transition. Embryos were injected with a combination of the following constructs: dUAS:Cherry-Wee1;CA-Cdc25 and dUAS:p120-mutFF;AnillinGFP. Movies were taken over 4 hours. Frames were extracted from a representative movie at indicated times from the tailbud stage. Scale bars, 25 *μ*m.

What mediates coupling of the cytokinetic machinery with the junctions? A recent study on the regulation of cytokinetic ring assembly identified p120-catenin, a component of the AJs, as a scaffold that restricts RhoA activation zone to the constricting ring^49^. This prompted us to hypothesise that p120-catenin, a well-known target of the Src kinase^50^, could be the factor that delocalises Anillin to the junctions in the absence of cues coming from the mitotic spindle. To test this hypothesis, we attempted to modulate the p120-catenin function in extruding vSrc cells. Two tyrosine residues Y217 and Y228 of p120-catenin when phosphorylated by Src are known to promote the interaction between the Src kinase and RhoA^51^. Therefore, we created a phosphomimetic mutant versions of p120-catenin in which these tyrosine residues were replaced with phenylalanines (FF) to mimic lack of phosphorylation (Fig. 7B). Interestingly, expression of the p120-catenin FF mutant together with vSrc already significantly attenuated extrusion, presumably acting as a dominant negative form in this context (Fig. 7C). This suggests that vSrc modulates adherens junctions to couple with the cytokinetic machinery.

We then sought if modified p120-catenin could link the cytokinetic ring to the junctions in a cell cycle-dependent manner, mimicking vSrc-like activity during cell extrusion. Expression of the p120-catenin EE mutant, in which the Src-dependent phosphorylation sites are replaced with glutamic acids to mimic a permanent state of phosphorylation, with concomitant expression of the Wee1 kinase and CA-Cdc25 phosphatase resulted in a G2/M arrest phenotype in normal EVL. In the presence of these three factors, we indeed observed Anillin-GFP recruited to the junctions. In some cases, the cells expressing these factors underwent basal extrusion accompanied by immediate cell death (Fig. 7D, Movie 9). Importantly, without the EE mutant of p120-catenin, expression of Wee1 and CA-Cdc25 was not sufficient to recruit Anillin-GFP to the junctions in cells arrested at the G2/M transition (Fig. 7E). In rare cases of basal extrusion due to protein overexpression in these embryos, Anillin was not recruited to the junctions and this type of extrusion appeared to be Anillin-ring independent (Fig. S7B). Finally, when p120-mutant-FF, instead of p120-mutant-EE, was expressed alongside Anillin-GFP in cells arrested at the G2/M transition, Anillin could not be stably recruited to the junctions, form a ring or facilitate cell extrusion (Fig. 7F). This last observation proved that phosphorylation of p120-catenin by vSrc on residues Y217 and Y228 was indeed responsible for the recruitment of Anillin to the junctions and drove the apicobasal split of vSrc cell during extrusion.

Collectively, Src activation in the EVL leads to altered cell cycle progression, assembly of a contractile ring initially parallel to the surface of the embryo through AJs in the prolonged G2 phase of the cell cycle and extrusion via constriction of this ring in early mitosis. During extrusion, the misoriented ring constricts and separates the basal from the apical part of the cell releasing both from the epithelium.

### vSrc promotes apical polarity shift and survival to enable extrusion

So far, we have identified crucial cellular process and property that need to be modified by the vSrc kinase for extrusion to occur: the cell cycle and AJs. However, when we tried to mimic vSrc-like changes in cells without the active kinase, extrusion occurred only occasionally, was basal instead of apical, and was associated with cell death (Fig. 7D). Hence, we wondered whether modulating cell polarity downstream of Src activation could lead to a change in directionality of extrusion. It has been shown that Src fine-tunes the activity of the small GTPase Cdc42 both directly and indirectly downstream of EGF stimulation^52^. Cdc42 has pivotal roles in establishing apicobasal polarity in all eukaryotic cells^53,54^ and in regulating the apical polarity complex aPKC-Par3-Par6 in a manner conserved among different species^55–57^. Therefore, Cdc42 could be a good candidate to link Src with polarity. When a dominant negative form of a downstream mediator of Cdc42, atypical protein kinase C (DN-aPKC), which contains only the N-terminal regulatory domain targeted to the membrane^58^, was expressed together with vSrc, it inhibited vSrc-driven extrusion (Fig. 8A). This suggests a role for the modulation of apicobasal polarity in vSrc-mediated extrusion.

Apart from modulating the cell cycle, AJs and cell polarity, we speculated that Src activation involves promoting cell survival, as demonstrated previously^59^. To reconstitute vSrc-like cell extrusion, finally we expressed all the components: the cell cycle modulators Wee1 and CA-Cdc25, the AJs’ component recruiting Anillin p120-mutant-EE, the polarity modulator constitutively active membrane-bound aPKC (myr-aPKC) and the pro-survival protein XIAP together in EVL cells. Indeed, combining all the components that mimic Src activation in the EVL resulted in vSrc-like extrusion: apicobasal split, where extruded cells did not die immediately (Fig. 8B, C, Movies 10 and 11). Thus, we managed to pinpoint four effector pathways downstream of vSrc that coordinate apicobasal extrusion: cell cycle via modulating Cdk1 and Pp2a, AJs via p120 catenin, apicobasal polarity and cell survival (Fig. 8D, S8C).

**Fig. 8.**
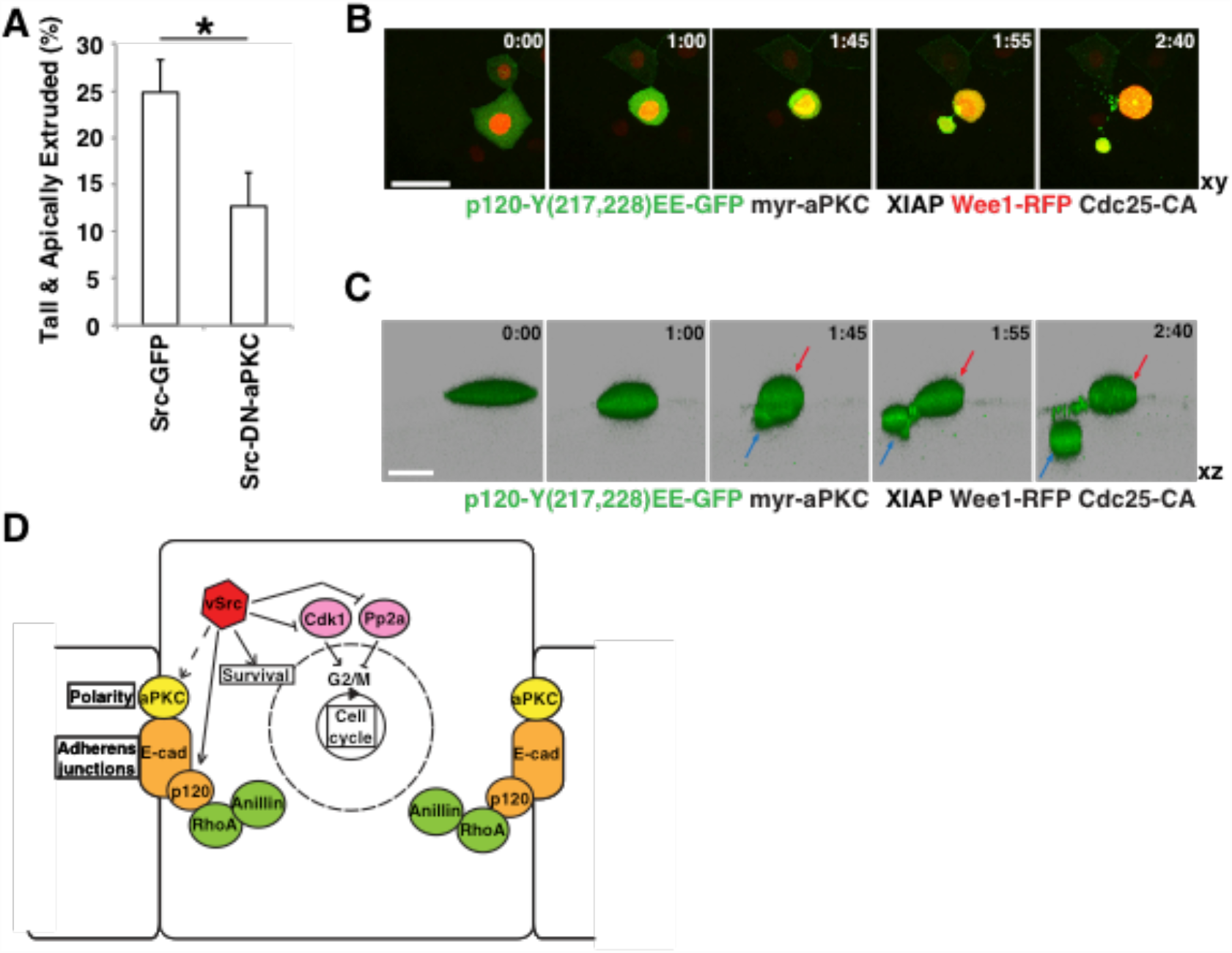
vSrc-like extrusion can be reproduced by modulating the cell cycle, junctions and polarity in a vSrc-like manner. **(A)** The effect of dominant negative aPKC on vSrc-driven extrusion. Embryos were injected with the following constructs: dUAS:EGFP-vSrc and dUAS:EGFP-vSrc;DN-aPKC. Data are mean ± s.d. 3 independent experiments (total number of embryos: n_Src_ = 31; n_Src,DN-aPKC_ = 31). *P < 0.05. **(B)** Time-lapse imaging of vSrc-like extrusion induced by coexpression of p120-mutEE, myr-aPKC and the apoptotic inhibitor XIAP in G2/M-arrested cells. Embryos were injected with a combination of the following constructs: dUAS:Cherry-Wee1;CA-Cdc25, dUAS:p120-mutFF;myr-aPKC and Krt18:XIAP. Movies were taken over 4 hours. Frames were extracted from a representative movie at indicated times from the tailbud stage. Scale bars, 50 *μ*m. **(C)** Time-lapse imaging of vSrc-like cell extrusion in (B) segmented using the Imaris software. The surface function was used to segment GFP positive cells over time. In this cross section of the embryo (xz view), a cell is undergoing an apicobasal split (apical part is marked with red arrows and the basal part with blue arrows). Scale bars, 25 *μ*m. **(D)** A schematic model of vSrc-driven cell extrusion. vSrc interferes with the cell cycle, and modulates adherens junctions, cell survival and apicobasal polarity, leading to apicobasal extrusion. vSrc-expressing cell becomes taller than its neighbours. Cell cycle regulators are hijacked; Pp2A is inactivated earlier in the cell cycle, but counteracting Cdk1 inhibition results in G2/M arrest instead of mitosis. The nuclear envelope becomes partially permeable and Anillin is recruited to the adherens junctions by modified p120-catenin, presumably through active RhoA. A contractile junctional ring assembles parallel to the plane of the epithelium, constricts in early mitosis and releases the cell from the epithelium. vSrc-mediated modulation of the apicobasal polarity complex (e.g. aPKC) promotes the predominantly apical direction of extrusion. Immediate cell death is avoided due to vSrc promoting cell survival.

## Discussion

In this report we have used the early zebrafish embryo to study oncogenic extrusion primarily based on high-resolution live imaging. We found that vSrc-mediated apicobasal extrusion is executed by hijacking the cell cycle and rewiring cytokinesis. vSrc drives EVL cells into the G2 phase of the cell cycle and initially blocks further progress. During this period, Src activation leads to the reorganisation of the zonula adherens (ZA) through incorporation of Anillin recruited by vSrc-modified p120-catenin, a component of AJs. With a contractile junctional ring assembled, the cell enters mitosis and the ring constricts, facilitating apicobasal extrusion of the vSrc cell. During extrusion, the larger part of the cell containing the nucleus is released apically. Thus, premature and rewired cytokinesis occurs in early prophase before NEB or mitotic spindle assembly.

Such premature cytokinesis has been described in unfertilised syncytial eggs in *Drosophila*, which physiologically remain in the M-phase^60^, despite the fact that there are no microtubule bundles present in these eggs outside of a small peripheral meiotic spindle. When injected locally with CDK1 inhibitors or an active RhoA, the embryo forms *de novo* a premature contractile structure that resembles a cytokinetic ring with Actin, Myosin and Anillin at the site of injection. In our system in the absence of a mitotic spindle, vSrc appears to generate a narrow zone of active RhoA by modulating p120-catenin^49–51^. This modulation results in recruitment of the cytokinetic scaffold Anillin and assembly of a premature misoriented cytokinetic ring by rebuilding the already existing junctional Actomyosin ring. To resolve the dual roles of RhoA in mediating cytokinesis and junctional integrity in extruding vSrc cells, optogenetic approaches will be necessary in the future^61^.

Transformation of a junctional Actomyosin ring into a contractile ring has been described previously in the context of apoptotic extrusion, where the ring was formed in the neighbours^62^. The process relies on the reorganisation of short into co-aligned perijunctional Actin bundles. This rearrangement is mediated by the Actin-binding protein Coronin B1 which is recruited to AJs by E-cadherin. Coronin B1 is not only required to assemble ZA in epithelial cells, but also to rearrange Actomyosin in the neighbours upon cell death. When a cell dies within an epithelium, two junctional Actomyosin rings can be seen at the level of AJs: one in the dying cell and the other in the neighbours. The neighbouring ring becomes thicker and relocalises basally to facilitate apical extrusion, while the ring in the dying cell remains stationary. In contrast, during vSrc cell extrusion, we observe that the autonomous ring changes in structure, becomes contractile and relocalises from the junctions to an oblique position. Despite being fundamentally different in terms of the mechanisms for generating force either autonomously or non-autonomously, these two processes are also remarkably similar. Both rely on modification of the ZA into a fully contractile ring to facilitate extrusion, and appear to be regulated by RhoA^63^. While the Actomyosin ring in neighbours of the dying cell is controlled by p115RhoGEF^64^, nothing is known about regulators of the autonomous contractile ring, which remain to be identified in the future.

Another issue to consider is the non-cell-autonomous contribution to apoptotic and oncogenic cell extrusion. Micheal et al.^62^ show that assembly of a contractile ring in neighbouring cells depends on apoptotic shrinkage of the dying cell, pulling on the neighbouring junctions and mechanosensing through E-cadherin. In case of vSrc-induced extrusion, a cell-autonomous active contraction via the pseudo cytokinetic ring generates a force that presumably pulls at the AJs of the neighbours. In this scenario, the direction of the forces that act on neighbouring AJs in dead and oncogenic extrusion is the same, but the strength would likely differ. Together, these findings raise the intriguing possibility that the pulling force on E-cadherin determines the mechanism used for extrusion by the neighbouring cells. Importantly, there is a difference in the non-cell-autonomous response. The Actin ring in the neighbours is less pronounced in Src-driven extrusion than in apoptotic extrusion^8,63^. Instead, neighbours of transformed cells appear to employ the Actin cross-liking protein Filamin to facilitate this process^3^. Remarkably, non-autonomous recruitment of both Actin and Filamin is regulated by RhoA^5,63^. Since p115RhoGEF regulates RhoA in the neighbours of a dying cell and mediates Actomyosin ring assembly^64^, another RhoA GEFs/GAPs could be involved in the regulation of RhoA and Filamin in extrusion of transformed cells. In future studies, it will be crucial to clarify whether E-cadherin signalling that acts as part of a mechanosensor could be upstream of differential RhoA activation in the neighbours, and to separate regulators of RhoA upstream of either Actin or Filamin in each of these processes.

It appears that RhoA GEFs/GAPs in transformed cells and their neighbours are also key in controlling the direction of extrusion: apical - outside of the embryo or basal - towards the deep cells of the embryo. It has been shown that p115RhoGEF plays a crucial role in determining where the Actomyosin ring is assembled in the neighbours of a dying cell^64^. Moreover, a recent study in oncogenic extrusion in the *Drosophila* wing imaginal disk implicates RhoGEF2, a fly homolog of p115RhoGEF, in determining directionality^65^. The presence of RhoGEF2 is linked to “tumour hotspots” with predominant apical extrusion whereas “tumour coldspots” is associated mostly with basal extrusion. However, our results (Figs. 8B,C) show that autonomous polarity change is sufficient to reverse the direction of extrusion. These seemingly opposing observations about whether directionality of extrusion is controlled cell-autonomously or non-cell-autonomously may be consolidated by the hypothesis that positioning of the ZA in the neighbours could be responsible for driving extrusion^62^. Since AJs connect transformed cells to their neighbours and give rise to the Actomyosin ring facilitating extrusion, positioning of the AJs should be crucial to determine the direction of extrusion and could in theory be regulated from both sides: the extruding cell and its neighbours.

Our data reveal that autonomous regulation of apicobasal polarity is necessary for vSrc-mediated extrusion (Fig. 8A) and may contribute to regulating its directionality. Recent findings on apical domain expansion in epithelial cells shed light on how this process could be involved in extrusion^57,66^. Dbl3 is a regulatory GEF of Cdc42, which has been shown to be necessary for oncogenic extusion^7^. Dbl3 is responsible for apical localisation and activation of Cdc42 and for expansion of the apical domain in epithelial cells through the regulation of the apical polarity complex aPKC-Par3-Par6. Downstream of Cdc42, the myosin kinase MRCK promotes myosin flow that separates apical aPKC-Par6 from junctional Par3, a step crucial for epithelial differentiation. Interestingly, MRCK was found differentially phosphorylated in H-Ras^V12^ cells upon interacting with normal cells prior to extrusion^6^. It appears that overexpression of Dbl3 promotes apical expansion resembling rounding up prior to extrusion^66^. Hence, this pathway may play a role in promoting rounding alongside RhoA in cell cycle-dependent extrusion or possibly on its own in cell cycle-independent extrusion. Further studies will be necessary to clarify this point.

During proliferation and crowding-induced extrusion, normal epithelia have an intrinsic mechanism of regulating their density, which dictates whether extrusion or division occurs. Preferential divisions occur in low-density epithelial sites (stretch), while crowding (squeeze) induces extrusion. This process is mediated by the stretch-activated calcium channel Piezo1^67^. However, it remains unclear whether this mechanism also functions in some squamous epithelia, such as the EVL, where all the cells are constantly stretched encasing the yolk and the deep cells. Even less is known as to whether Piezo1 mediates extrusion of transformed cells. Since vSrc-transformed cells become extruded instead of dividing, does that mean they somehow imitate crowding? We suspect that Piezo1 may not be involved in this type of extrusion, as our results indicate that the process of vSrc – induced extrusion is primarily driven autonomously, thereby resulting in pulling of the neighbours rather than pushing, which is apparently different to Piezo1-mediated extrusion. In our view, extrusion of transformed cells resembles cell death-induced extrusion and, as mentioned above, may employ some of the mechanisms involved in this process, in regards to the response in the neighbours. Nevertheless, it remains to be elucidated whether Piezo1 mediates extrusion of transformed cells.

Finally, it will be worth investigating whether cell extrusion induced by other oncogenes occurs in a cell cycle-dependent manner. If the mechanism is similar, blocking proliferation with drugs while treating carcinogenesis may impair the primary EDAC response and should be reconsidered.

Overall, our study uncovers a novel mechanism underlying EDAC. Further investigation will allow us to identify regulators of GTPases (in particular, RhoA) that regulate different aspects of extrusion in both cell-autonomous and cell-non-autonomous mechanisms. Understanding the coordination of timing, apical polarity and junctional integrity may eventually lead to potential therapies to boost EDAC.

## Methods

### Generation and maintenance of transgenic fish lines

The maintenance of fish and the collection of embryos were performed as described before^68^. The line Tg(Krt18:KalTA4-ERT2) was previously established^3,10^. To establish Tg(Krt18:Lifeact-Ruby) and Tg(Krt18:CcnB1-GFP) lines we used the vector pBR-Tol2-Krt18 generated previously^8^ and transferred Lifeact-Ruby^69^ and CcnB1-GFP (see Meterials), respectively, downstream of the Krt18 promoter. The resulting constructs (30 pg) were then coinjected with Tol2 RNA (7.5 pg) in the morpholino buffer (5 mM HEPES pH 7.5, 200 mM KCl) into one-cell wild type embryos. The embryos positive for RFP (Lifeact-Ruby) and GFP (CcnB1-GFP) expression at 10 hours post-fertilization (hpf) were raised to adulthood, and crossed with wild type fish to identify founder fish. Embryos from potential founders were imaged to select the optimal level of expression at which no overexpression phenotype could be observed. The founder fish were out-crossed with WT, and the F1 fish were selected on the basis of their fluorescent signal. All the embryos for experiments were obtained from crossing fish heterozygous with the Tg(Krt18:KalTA4-ERT2) line.

### Microinjection and confocal imaging of zebrafish embryos

Embryos were injected with a single construct (16–20 pg) or multiple constructs (combined amount of DNA was 20 pg) and Tol2 RNA (5 pg) in the morpholino buffer (5 mM HEPES pH 7.5, 200 mM KCl) into the cell at one-cell stage, and treated with 0.5 mM 4-hydroxy tamoxifen (Sigma H7904, a stock of 5mM in ethanol) at 50–70% epiboly as described^2^. For live imaging, after 2 hours of treatment, embryos were mounted in 0.8% low-melting agarose in fish water prior to confocal analysis. For immunofluorescence and quantification of extrusion rates, they were fixed in 4% PFA/PBS at 2.5–3 h after induction, stained and mounted in 1% low-melting agarose in PBS prior to confocal analyses. Movies were taken over 4 hours or over 8 hours (cell cycle analysis with the Krt18:CcnB1-GFP line). Confocal images were taken using a 25x 0.95 NA water-immersion lens on a high-resolution single photon microscope Leica TCS SP8 and were analysed using the Imaris software (Bitplane).

### Immunostaining of fish embryos

At 10 hpf GFP- or RFP-positive embryos were selected and dechorionated in 1% agarose plates to avoid damage. Embryos were fixed in a fresh solution of 4% PFA/PBS overnight at 4^o^C and subsequently washed 3x in PBS. Permeabilisation was performed for 15 min in PBS/0.5%TritonX-100 (PBSTr). Blocking in PBSTr/10% Goat serum/1% DMSO (Blocking buffer) lasted >1 h. Embryos were incubated with 1^st^ antibody in 200 *μ*l in Blocking buffer @ 4°C O/N, then washed with PBSTr 3–6x for 30 min in total. Incubation with 2^nd^ antibody in 200 *μ*l Blocking buffer lasted 3–4 hours at room temperature, followed by washes with PBSTr 3–6x for 30 min in total. Phalloidin staining was performed for 30 min in PBSTr/ 10% Goat serum.

### Cell culture experiments

MDCK cell lines were used in this study. The parental MDCK cells were a gift from Walter Birchmeier. MDCK and MDCK-pTR cSrc-Y527F-GFP lines were cultured as previously described^7,70^. To establish MDCK-pTR cSrc-Y527F-GFP line stably expressing FUCCI cell cycle markers mCherry-hCdt1(30/120) and mTurquoise-hGem(1/110), MDCK-pTR-cSrc-Y527F-GFP cells were transfected with P2A Fucci2.2_pCSII-CMV vector (a kind gift from Dr. Miyawaki) together with a pcDNA3.1 as a selection vector using Lipofectamine 2000 (Life Technologies), followed by selection in the medium containing 800 μg/mL of G418 (Gibco), 5 μg/mL of blasticidin, and 400 μg/mL of zeocin. To induce Src-expression, 2 μg ml^−1^ of tetracycline (Sigma-Aldrich) was added to the medium. For immunofluorescence and time-lapse experiments, cells were cultured on type-I collagen gels from Nitta Gelatin (Nitta Cellmatrix type 1-A; Osaka, Japan) as previously described^7^. For immunofluorescence, mixed cultures of cells (MDCK: Src = 50: 1) were plated and incubated for 8 hours, before adding tetracycline. To avoid differences in cell density which could affect extrusion rates, proliferation inhibitors were added to the medium 16 hours after tetracycline at following concentrations: hydroxyurea (2 mM) or Ro-3306 (10 *μ*M). Cells were fixed and stained as described previously^6^. Immunofluorescence images were taken with the Olympus FV1000 or FV1200 system and Olympus FV10-ASW software. Images were analysed with MetaMorph software (Universal imaging). For time-lapse imaging, following a 4 hour-tetracycline treatment, small groups of GFP-positive cells were chosen for imaging with Olympus IX81-ZDC”(Olympus) and images were taken and analysed with Metamorph software (Molecular Devices). For Western blotting cells were plated in plastic dishes and induced with tetracycline for 8 hours before lysis. Western blotting was carried out as previously described^71^. Primary antibodies were used at 1:1000. The western blotting data were analysed using ImageJ (NIH). For FACS analysis, MDCK cells were incubated with or without proliferation inhibitors as before for 16 hours prior to staining with Hoechst 33342 dye (1 ug/uL; ThermoFisher Scientific). After trypsinisation and straining, cells were counted, resuspended in 2% FBS/PBS, stained with propidium iodide and analysed for DNA content using FACSAriaTM II (BD Biosciences).

### Data analysis

For data analyses, two-tailed Student’s t-tests were used to determine P-values. P-values less than 0.05 were considered to be significant. Extrusion rates in fixed embryos were expressed as the number of “extruded” and “tall” cells (unless indicated otherwise) by the total number of GFP- or RFP-positive cells in the embryo. As “extruded” we classified cells that are no longer a part of the monolayer (their junctions closed off or nearly closed off up to 90%). As “tall” we classified cells that were at least double the height of an average EVL cell, displaying signs of early extrusion, such as rounding. Only embryos with between 5–50 GFP- or RFP-positive cells were taken into consideration.

Proliferation rates in living embryos were expressed as the number of divisions over 4 hours by the total number of cells at the beginning of the movie. Only embryos with between 5–35 GFP- or RFP-positive cells were taken into consideration. To measure chromatin volume, H2B-GFP signal was used to segment the GFP-positive region in the cell undergoing division or extrusion over time using the surface function of Imaris software. A constant threshold was used to avoid bias between different movies. The moment of mitosis or extrusion was set as point 0 and volumes from different movies were aligned according to time before and after extrusion and averaged to create graph Fig. 3G. To measure the intensity of the CcnB1-GFP signal, segmentation was performed in the red channel on the basis of the signal from the cell surface marker myr-Cherry using the surface function of the Imaris software. The segmented cell surface was then used to calculate the average intensity of the green channel and cell volume (Fig. 4D, E, F). To define the position of the Anillin ring, Imaris spot function was used to determine points within the plane of the ring and the plane of the surface of the embryo. Extracted coordinates of the spots where then fed into a MATLab function (based on affine_fit(X)) to calculate the angle between two planes (Fig. 2E).

### Data availability statement

The datasets generated and/or analysed during the current study are available from the corresponding author on reasonable request.

### Code availability statement

MatLab code used to quantify the angle of the Anillin ring will be released upon publication.

## Acknowledgements

We thank members of the Wilson, Rihel, Bianco and Tada labs for help, advice and sharing reagents, and the UCL fish facility for excellent zebrafish care. We would like to thank David Whitmore, David Kimelman, Sergei Sokol, Marina Mione, Jon Clarke, Luccia Poggi, Roberto Mayor, Caroline Brennan for constructs they provided. We especially would like to thank Buzz Baum, Karl Matter, Jon Clarke, Richard Poole and Snezhka Oliferenko for critical reading of the manuscript and advice. This work is funded by the Cancer Research UK. YF was supported by Japan Society for the Promotion of Science (JSPS) Grant-in-Aid for Scientific Research on Innovative Areas 26114001, Grant-in-Aid for Scientific Research (A) 26250026, the Naito Foundation, and the Takeda Science Foundation.

## Author contributions

K.A. designed experiments, generated and analysed most of the data. M.K. generated MDCK-FUCCI line. K.A., M.K. and R.N. performed live-imaging in MDCK cells. R.N. performed Western blots in MDCK cells. M.T. conceived and designed the fish model system and generated fish lines. Y.F. conceived and designed the MDCK model system. K.A. and M.T. conceived and designed the study. The manuscript was written by K.A. and M.T. with assistance from the other authors.

## Materials

### Constructs

All the constructs used for experiments were based on the pBR-Tol2 vector with either Krt18 promoter or the 5xUAS element driving expression in one (UAS, Krt18) or both (dUAS, dKrt18) directions^3,8,10^. To generate the dKrt18 vector, we placed the endogenous basal promoter (-150 bp from the transcription staring site of *krt18* gene) to –5kb upstream of the EVL-regulatory sequence of the Krt18 promoter in the reverse complementary orientation (see Fig S1). Previously published constructs used in this work were: dUAS:EGFP-vSrc^10^, UAS:EGFP-vSrc^3,10^, UAS:myr-Cherry-vSrc^3^. On the basis of the four basic pBR-Tol2 vectors with single or double promoters we created a number of new constructs using the InFusion system (Clontech). To make new constructs in most cases we used cDNA from zebrafish embryos unless otherwise indicated. The following previously cloned cDNAs were gifts: Wee1, p20 and p21 from David Whitmore^72,73^, CA-Cdc25 (Cdc25–3S/T-A) from David Kimelman^35^, aPKC (rat) from Sergei Sokol^74^, Dcx-GFP from Marina Mione^75^, H2B-GFP from Jon Clarke, Anillin from Luccia Poggi^26^, p120-wt (mouse) from Roberto Mayor^76^, DAPK1 from Caroline Brennan. The following cDNAs were cloned from a cDNA library created using 24-hour old zebrafish embryos: Pp2a (ZDB-GENE-050417–441), RhoA (ZDB-GENE-040426–2150), CcnB1 (ZDB-GENE-000406–10), Cdk1 (ZDB-GENE-010320–1), XIAP (ZDB-GENE-030825–7) based on the ZFIN database. Indicated point mutations and deletions were achieved using the InFusion method (Clontech) and confirmed by sequencing (Source BioScience). The specific created mutations were: Y307F in CA-Pp2a, Q63L in CA-RhoA, T19N in DN-RhoA, T14A and Y15F in CA-Cdk1, Δ(1–740) in DN-Anillin, Δ(201–591) in DN-aPKC, Y217,228F in p120-mutFF, Y217,228E in p120-mutEE. nucGFP was created by fusing 2xNLS Sv40 with NLS from Wee1 (RNNRKRSHWN), hmAzami-Green and EGFP. CDK1pep expressing construct was created by fusing FLAG, mKO2 fluorophore, CDK1 peptide (KIEKIGEGTYGVVYK) and 2xHA tag.

On the basis of the dUAS:EGFP-vSrc construct we created: dUAS:EGFP-vSrc;Wee1, dUAS:EGFP-vSrc;CA-Cdc25, dUAS:EGFP-vSrc;p20, dUAS:EGFP-vSrc;p21, dUAS:EGFP-vSrc;Pp2a-Y307F, dUAS:EGFP-vSrc;CA-RhoA, dUAS:EGFP-vSrc;DN-RhoA and dUAS:EGFP-vSrc;DN-aPKC. We also replaced the EGFP in the dUAS:EGFP-vSrc construct with myr-Cherry to obtain dUAS:myr-Cherry-vSrc and subsequently used it to make the following constructs: dUAS:myr-Cherry-vSrc;XIAP, dUAS:myr-Cherry-vSrc;Dcx-GFP, dUAS:myr-Cherry-vSrc;H2B-GFP, dUAS:myr-Cherry-vSrc;nucGFP, dUAS:myr-Cherry-vSrc;Anillin-GFP, dUAS:myr-Cherry-vSrc;DN-Anillin-GFP, dUAS:myr-Cherry-vSrc;p120-wt, dUAS:myr-Cherry-vSrc;p120-mutFF. We used the original pBR-Tol2-dUAS vector to create the following constructs: dUAS:Cherry-Wee1;CA-Cdc25, dUAS:p120-mutEE;AnillinGFP, dUAS:myr-Cherry;AnillinGFP, dUAS:p120-mutFF;AnillinGFP, dUAS:p120-mutFF;myr-aPKC, dUAS:myr-Cherry;GFP-Emerin, dUAS:Dcx-GFP;H2B-RFP, dUAS:myr-Cherry;H2B-GFP, dUAS:myr-Cherry;Anillin-GFP, dUAS:GFP-CAAX;CA-RhoA, dUAS:myr-Cherry;DAPK1. We used the pBR-Tol2-UAS to create: UAS:myr-Cherry. We used pBR-Tol2-Krt18 to create the following constructs: Krt18:XIAP, Krt18:CcnB1-GFP. We used the pBR-Tol2-dKrt18 to create the following constructs: dKrt18:H2B-GFP;myr-Cherry, dKrt18:myr-Cherry, dKrt18:Cherry-Wee1 and dKrt18:Cherry-Wee1;CA-Cdk1.

### Antibodies, morpholinos and inhibitors

Anti-GFP antibody was from Abcam (13970). Anti-RFP antibody was from MBL (PM005). Anti-phospho-CDK1 Tyr15 antibody was from Cell Signaling (4539). Anti-phospho-MLC2 Thr18/Ser19 antibody was from Cell Signaling (3674). Anti-phospho-Histone H3 Ser10 antibody was from Upstate (MERCK: 06–570). Anti-active Caspase 3 antibody was from BD Biosciences (559565). Secondary antibodies were from Invitrogen Molecular Probes. Phalloidin-Atto 647N was from Sigma.

For knockdown experiments in zebrafish, we used Emi1 MO^77^, a gift from Jon Clarke. 1 nL of 0.5 mM morpholino (Emi1 MO or control MO) solution was injected into the yolk following a DNA injection. For chemical inhibition of proliferation, we used aphidocholin (150 *μ*M) and hydroxyurea (20 mM) from Sigma in fish water containing 4% DMSO. Inhibitors were added together with tamoxifen, at 50–70% epiboly. To generate mitotic spindle defects, Eg5 (Kif11) inhibitor STLC from Alfa Aesar was used at 0.874 mM added together with tamoxifen, at 50–70% epiboly and during imaging.

